# A chemical screen identifies p38 MAPK inhibition as a candidate neuroprotective strategy for combinatorial SMA therapy

**DOI:** 10.1101/2025.03.06.641938

**Authors:** Maria J. Carlini, Jorge Espinoza-Derout, Meaghan Van Alstyne, Sarah Tisdale, Eileen Workman, Marina K. Triplett, Ivan Tattoli, Shubhi Yadav, Christopher E. Henderson, Martin D. Watterson, Livio Pellizzoni

## Abstract

Spinal muscular atrophy (SMA) is a neurodegenerative disease caused by ubiquitous deficiency in the survival motor neuron (SMN) protein. The identification of effectors and modifiers of pathogenic events downstream of SMN deficiency is key to understanding disease mechanisms and broadening the range of targets for developing SMA therapies that can complement SMN upregulation. Here, we report a cell-based phenotypic screen for chemical modifiers of SMN biology that identified inhibitors of p38 mitogen-activated protein kinase (p38 MAPK) as suppressors of proliferation defects induced by SMN deficiency in mouse fibroblasts. We further show that SMN deficiency induces p38 MAPK activation and that pharmacological inhibition of this pathway improves motor function in SMA mice through SMN-independent neuroprotective effects. Using a highly optimized p38 MAPK inhibitor (MW150) and a specific paradigm of combinatorial treatment in SMA mice, we observed synergistic enhancement of the phenotypic benefit induced by either MW150 or an SMN-inducing drug alone. By promoting survival of motor neurons, pharmacological inhibition of p38 MAPK synergizes with SMN induction and enables enhanced synaptic rewiring of motor neurons within sensory-motor spinal circuits, resulting in increased motor function, weight gain, and survival of SMA mice. Together, our studies identify the p38 MAPK pathway as a therapeutic target and MW150 as a candidate pharmacological approach for SMN-independent neuroprotection with clinical relevance for combination therapy in SMA.

## Introduction

Spinal muscular atrophy (SMA) is an inherited neurodegenerative disease characterized by loss of motor neurons and skeletal muscle atrophy, leading to motor dysfunction, paralysis and eventually death if left untreated (1, 2). SMA is caused by ubiquitous reduction in the levels of the survival motor neuron (SMN) protein, which results from homozygous loss of the *SMN1* gene and preservation of the paralog *SMN2* gene (3). Due to a single nucleotide difference affecting the splicing of exon 7 (4), *SMN2* produces low levels of full-length functional SMN protein that cannot compensate for the loss of *SMN1* leading to SMA (5). Accordingly, the time of onset and the severity of the disease inversely correlate with *SMN2* copy number and SMN protein levels (1, 2).

The SMN protein has critical functions in RNA regulation, including but not limited to the assembly of small nuclear ribonucleoproteins (snRNPs) that carry out pre-mRNA splicing and 3’-end processing of histone mRNAs (6, 7), and increasing evidence from preclinical studies in animal models links RNA processing defects to SMA pathology (8, 9, 18, 10–17). Furthermore, although selective degeneration of motor neurons and skeletal muscle atrophy are disease hallmarks, an expanding body of evidence directly implicates dysfunction of multiple neuronal and non-neuronal elements of sensory-motor circuits in the etiology of SMA (13, 19–27).

In recent years, therapeutic approaches that increase SMN expression through antisense oligonucleotides or small molecules that target *SMN2* splicing, as well as SMN replacement by gene therapy, have been approved for the treatment of SMA patients (28–33). Despite evidence of clinical efficacy, it is widely appreciated that none of these therapies represent a cure for the disease (34–37). The response to therapy is variable and a significant proportion of patients show modest improvement, especially when treatment is delayed. Accordingly, incomplete correction of disease symptoms combined with variability in the clinical response to current treatments represent outstanding challenges in the SMA field.

To address currently unmet needs of SMA patients, the development of combination therapies that can enhance the clinical benefit of SMN-inducing drugs is critical (37). This calls for new approaches to target cellular factors and molecular pathways that are disrupted downstream of SMN deficiency. However, a current limitation is the availability of disease-relevant targets that could be modulated pharmacologically. This requires increased knowledge of disease mechanisms as well as identification and validation of novel, SMN-independent therapeutic approaches as viable entry points for subsequent clinical testing. To date, the most clinically advanced efforts are focused on targeting myostatin activation as a means to counteract muscle weakness and atrophy (38, 39). In contrast, there has been a paucity of approaches targeting neuroprotection and the only candidate molecule (olesoxime) that advanced to clinical testing failed to demonstrate benefit in SMA patients (40, 41). Of note, none of these approaches have been designed to target mechanistically established molecular defects induced by SMN deficiency in SMA.

Here we sought to address these outstanding issues through the discovery of novel cellular targets and preclinical validation of disease-modifying pharmacological approaches that are SMN-independent and ideally suited for combinatorial treatment of SMA. To do so, we carried out a cell-based screen for chemical modifiers of SMN biology and identified inhibitors of p38 mitogen-activated protein kinase (p38 MAPK) as suppressors of cellular phenotypes induced by SMN deficiency in cultured mammalian cells. Furthermore, we provide proof-of-concept for the therapeutic benefit of pharmacological inhibition of this signaling pathway in a mouse model of SMA - either alone or in combination with an SMN inducing drug - which occurs through SMN-independent neuroprotective mechanisms promoting motor neuron survival. Importantly, these studies employed an isoform selective p38αMAPK inhibitor (MW150) – an experimental therapeutic in early-stage trials for neurodegenerative and neuropsychiatric disorders optimized for safety and low risk for drug-drug interaction – that has the desired properties for use in combination therapy. Thus, our findings identify p38 MAPK as a disease-relevant cellular target and MW150 as a candidate small molecule for SMN-independent neuroprotection in SMA.

## Results

### A cell-based chemical screen identifies candidate modifiers of SMN biology

We previously established a cell model system that uses proliferation as phenotypic readout of functional levels of human SMN produced from the *SMN2* gene in NIH3T3 fibroblasts genetically engineered for doxycycline (Dox)-regulated, shRNA-mediated knockdown of endogenous mouse Smn (42). Addition of Dox to NIH3T3-SMN2/Smn_RNAi_ cells induces Smn depletion by RNAi and severe impairment of cell proliferation which is dependent on low levels of human SMN produced by the *SMN2* gene. We also established a cell-based assay in 96-well format for the direct measure of NIH3T3 cell number using nuclear staining with Hoechst followed by automated, whole-well imaging (42). Here, we used this system in a phenotypic screen for chemical compounds that increase cell number of Smn-deficient NIH3T3 as a means to identify candidate small molecule modulators of SMN biology.

The screening design included pre-treatment of NIH3T3-SMN2/Smn_RNAi_ cells with Dox for 5 days prior to seeding in 96-well plates, addition of chemical compounds, incubation for 5 additional days in the presence of Dox, and readout of cell number (Figure 1A). Several parameters of the cell-based assay were optimized for the chemical screen (see Materials and Methods) and validated to ensure accuracy and specificity using SMN-C3 - a compound that potently promotes SMN2 exon 7 splicing and was further developed into the FDA-approved Risdiplam (43, 44) - as a benchmarking compound. Accordingly, proliferation of Smn-deficient NIH3T3-SMN2/Smn_RNAi_ cells was strongly increased by SMN-C3 relative to vehicle using our cell-based assay (Figure 1B). Dose-response analysis of SMN-C3 indicated strong activity (>3-fold increase in cell number at the lowest maximal effective concentration) and high potency (EC50=210nM) as well as specificity because no effects were found on the proliferation of Smn-deficient NIH3T3-Smn_RNAi_ cells that do not contain the *SMN2* gene (Figure 1D).

**Figure 1.**
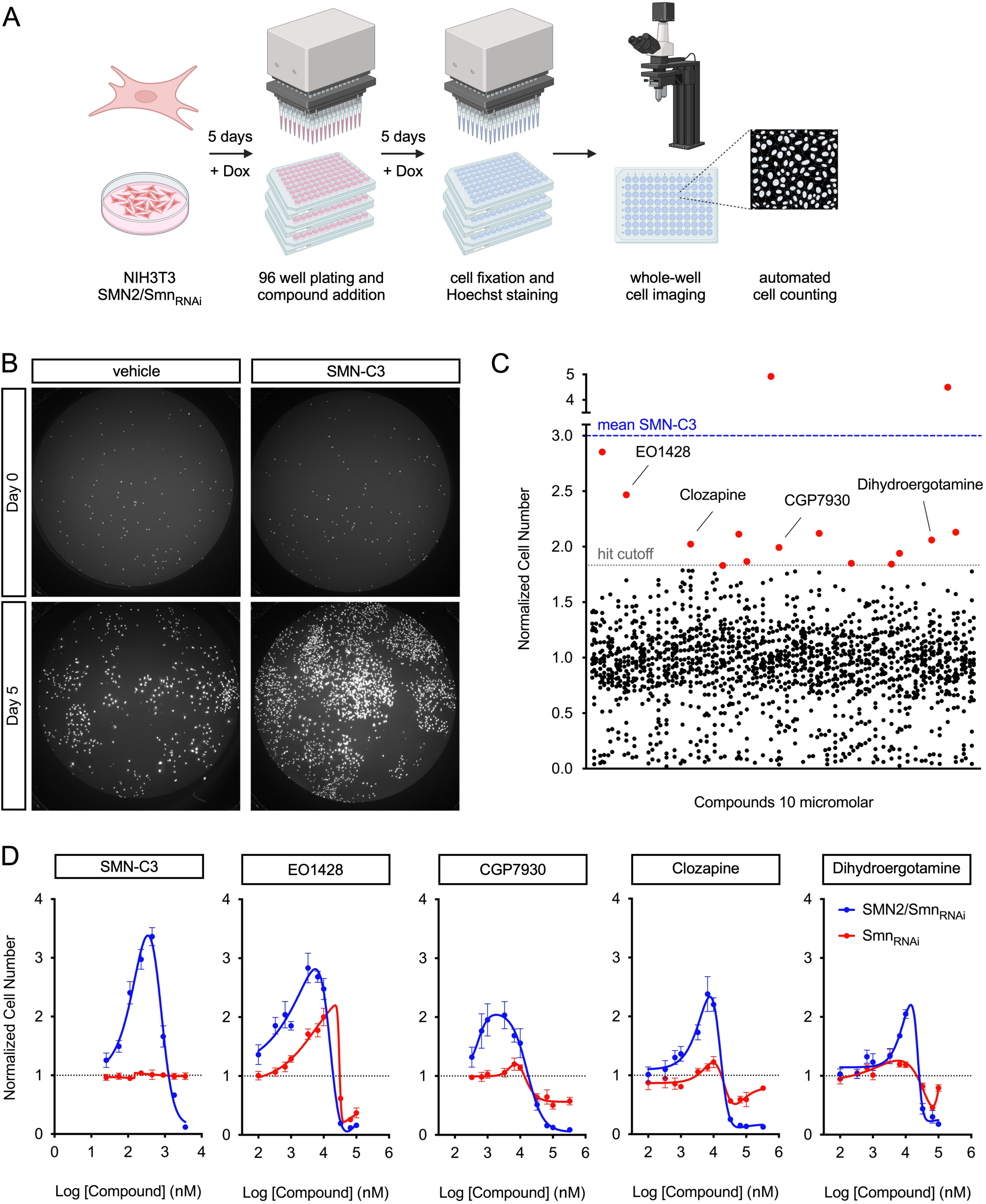
A cell-based chemical screen identifies candidate modifiers of SMN biology. (**A**) Schematic of the chemical screen design. NIH3T3-SMN2/Smn_RNAi_ cells were grown for 5 days in the presence of Dox prior to plating in 96-well format plates and addition of compounds from a chemical library. After additional 5 days in culture in the presence of Dox, cells were fixed and stained with Hoechst followed by automated whole-well imaging and determination of cell number. (**B**) Representative whole-well images of Dox-treated NIH3T3-SMN2/Smn_RNAi_ cells cultured in 96-well format in the presence of vehicle (DMSO) or SMN-C3 (500nM) for 4 hours (Day 0) or 5 days (Day 5) followed by fixation and nuclear staining with Hoechst. (**C**) Graph summarizing the results of the chemical screen. Each point represents the effect of a single compound on the proliferation of NIH3T3-SMN2/Smn_RNAi_ cells at day 5. The mean normalized cell number relative to vehicle treated cells from 3 replicates is shown. The mean of the benchmark compound (SMN-C3) and the hit cutoff are indicated by dotted lines. Primary hits are shown in red and the 4 compounds that passed secondary validation are labeled by name. (**D**) Dose-response analysis of the effects of the indicated compounds on cell proliferation in Dox-treated SMN2/Smn_RNAi_ and Smn_RNAi_ NIH3T3 cells. The graphs show mean and SEM (n=6) of normalized cell number relative to vehicle treated cells at day 5 for each tested concentration, and non-linear curve fitting.

Using the cell-based assay and design described above, we screened 1,307 chemical compounds with known biological activity. Dox-treated NIH3T3-SMN2/Smn_RNAi_ cells were incubated with a single compound per well at a concentration of 10 μM in the presence of Dox for 5 days followed by cell number determination. The screen was carried out in triplicate and hits were defined as compounds increasing cell number beyond the vehicle-treated mean plus 3 standard deviations (Figure 1C). According to these criteria, the screen yielded 15 compounds as initial hits, 4 of which (EO1428, CGP7930, Clozapine, and Dihydroergotamine mesylate) were confirmed after subsequent validation including counter-screen assays for specificity and dose-response experiments (Figure 1C and 1D, see also Materials and Methods). The validated compounds increased the proliferation of Smn-deficient NIH3T3-SMN2_low_/Smn_RNAi_ cells in a dose-dependent manner across a range of concentrations and had EC50 values of <10μM (Figure 1D), whereas they showed progressive toxicity at concentrations in the hundred micromolar range. Dose-response studies were also performed in Dox-treated NIH3T3-Smn_RNAi_ cells for a first insight into potential mechanisms of action, and EO1428 was also the only compound found to promote proliferation in these cells that do not contain the human *SMN2* gene (Figure 1D). Lastly, Western blot analysis demonstrated that none of the validated compounds act by increasing the low levels of SMN expression in Dox-treated NIH3T3-SMN2_low_/Smn_RNAi_ cells (Figure 2A). Thus, our cell-based screen identified a set of chemical modifiers of cellular proliferation defects induced by SMN deficiency.

**Figure 2.**
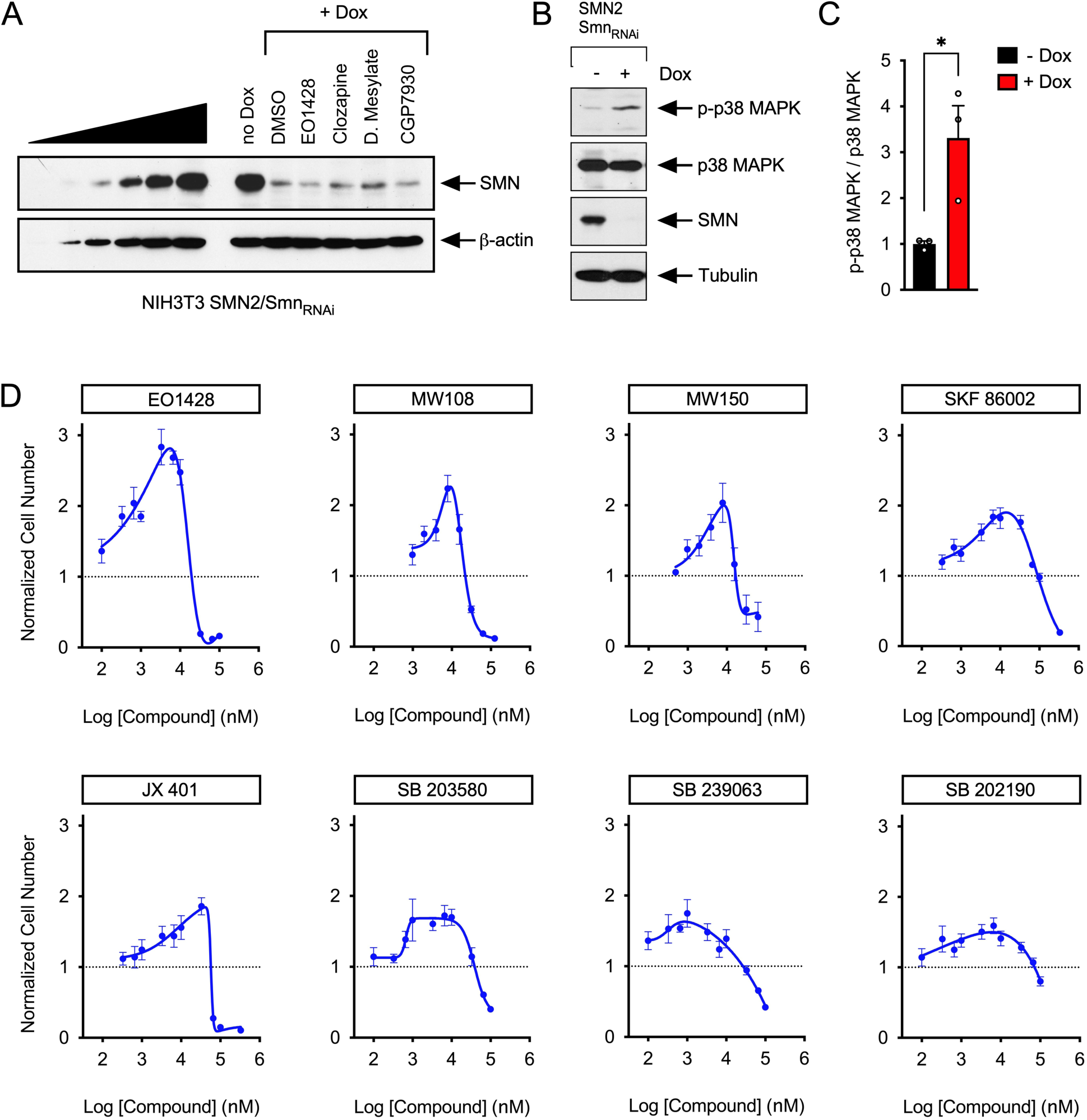
p38 MAPK inhibition promotes proliferation of SMN-deficient NIH3T3 cells. (**A**) Western blot analysis of SMN expression in untreated and Dox-treated NIH3T3-SMN2/Smn_RNAi_ cells cultured for 5 days in the presence of either DMSO or EO1428, Clozapine, Dihydroergotamine mesylate and CGP7930 at a concentration of 10μM. β-actin was used as a loading control. A two-fold serial dilution of extracts from untreated NIH3T3-SMN2/Smn_RNAi_ cells is shown on the left. (**B**) Western blot analysis of total and phosphorylated p38 MAPK in untreated and Dox-treated NIH3T3-SMN2/Smn_RNAi_ cells. SMN and tubulin were used as controls. (**C**) Quantification of the levels of phosphorylated p38 MAPK relative to total p38 MAPK in NIH3T3-SMN2/Smn_RNAi_ cells cultured with or without Dox from experiments as in (B). Normalized mean, SEM, and individual values from 3 independent biological replicates are shown. Two-tailed unpaired Student’s t-test. *P < 0.05. (**D**) Dose-response analysis of the effects of the indicated p38 MAPK inhibitors on the proliferation of Dox-treated SMN2/Smn_RNAi_ cells at day 5. The graphs show mean and SEM of normalized cell number relative to vehicle treated cells at day 5 (n=6) for each tested concentration, and non-linear curve fitting.

### p38 MAPK inhibitors increase proliferation of SMN deficient fibroblasts

The identification of EO1428 as the top validated hit in our phenotypic screening pointed to p38 MAPK inhibition as a candidate modifier of SMN deficiency. To investigate this further, we first looked at phosphorylation of Thr180/Tyr182 in p38 MAPK as an established marker of its activation (45). Western blot analysis showed a significant increase in the levels of phosphorylated p38 MAPK in Dox-treated, Smn-deficient NIH3T3-SMN2/Smn_RNAi_ cells relative to untreated controls (Figure 2B and 2C). Next, we performed dose-response studies of the effects of several chemically distinct p38 MAPK inhibitors, including the highly specific p38α MAPK inhibitors MW108 and MW150 (46–48), on the proliferation of SMN-deficient NIH3T3-SMN2/Smn_RNAi_ cells. Importantly, despite expected differences in potency and activity, all the inhibitors were effective in counteracting SMN-dependent cell proliferation deficits (Figure 2D). These results indicate that the cell proliferation phenotype induced by SMN deficiency in mouse fibroblasts is mediated, at least in part, by p38 MAPK activation and can be modified through pharmacological inhibition.

### SMN deficiency induces p38 MAPK activation in the spinal cord of SMA mice

To determine whether SMN deficiency induces p38 MAPK activation in a disease relevant context *in vivo*, we used the well-established SMNβ7 mouse model of SMA that has homozygous knockout of mouse Smn, contains two copies of the human *SMN2* gene, and is homozygous for the SMNΔ7 transgene (Smn^-/-^;SMN2^+/+^;SMNΔ7^+/+^) (49). Its pattern of motor unit loss closely matches that seen in untreated Type 1 SMA patients (50). We performed a longitudinal analysis of p38 MAPK activation in spinal cords isolated from WT and SMA mice at pre-symptomatic (P1) as well as early (P6), and late (P11) symptomatic stages of the disease by Western blot (Figure 3). This analysis showed a robust increase in the levels of phosphorylated p38 MAPK in the spinal cord of SMA mice relative to controls from P1 to P6 (Figure 3A-D), while no differences were found at P11 (Figure 3E and 3F). Thus, SMN deficiency induces the activation of p38 MAPK at the onset of SMA pathology in a severe mouse model of the disease.

**Figure 3.**
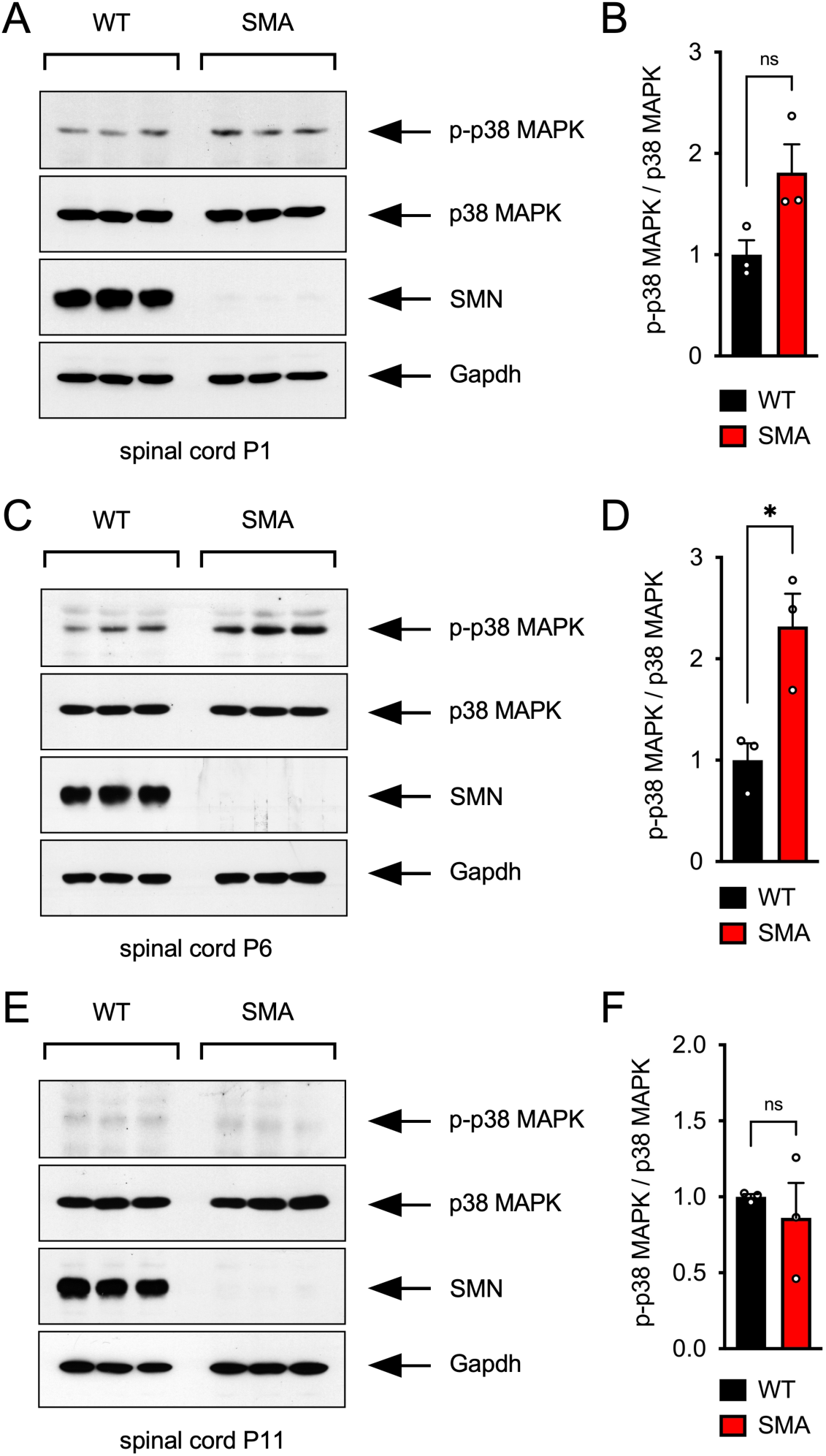
SMN deficiency induces p38 MAPK activation in the spinal cord of SMA mice. (**A**, **C**, **E**) Western blot analysis of total and phosphorylated p38 MAPK in the spinal cord of WT and SMA mice at P1 (A), P6 (C), and P11 (E). SMN and Gapdh were used as controls. (**B**, **D**, **F**) Quantification of the levels of phosphorylated p38 MAPK relative to total p38 MAPK from the experiments in A, C, and E. Normalized mean, SEM, and individual values from 3 independent biological replicates are shown. Two-tailed unpaired Student’s t-test. *P < 0.05; ns, not significant.

### p38 MAPK inhibition improves motor function of SMA mice through SMN-independent neuroprotective effects

To determine whether p38 MAPK activation contributes to SMA pathogenesis, we investigated the effects of its pharmacological inhibition in SMA mice. Most of the commercially available p38 MAPK inhibitors have the potential for off-target effects due to their mixed kinase inhibition activity beyond p38 MAPK, and limited information is available regarding their CNS penetrance. In contrast, MW150 is a potent and highly selective, blood brain barrier-permeable inhibitor of p38αMAPK that has previously been shown to be safe and effective in mouse models of Alzheimer’s disease (AD) and other CNS disorders (46–48, 51–54) and is currently under clinical development. Therefore, we selected MW150 for these *in vivo* studies.

First, we investigated the exposure levels of MW150 in plasma and brain of neonatal SMA mice following a single intraperitoneal (IP) injection of four escalating drug concentrations (2.5, 5, 10 and 20mg/kg) at P10. Brain and plasma samples were collected 3 hours after injection followed by analysis of MW150 concentration by LC-MS/MS. Both plasma and brain showed a dose-dependent linear increase in the levels of MW150 that was proportional to the amount of the drug injected in SMA mice (Supplementary Figure 1A and 1B). Moreover, a relatively constant brain-to-plasma ratio of ∼6 was found at all doses tested (Supplementary Figure 1C). These studies confirmed the outstanding brain permeability of MW150 and demonstrated that MW150 drug exposure levels known to effectively inhibit p38αMAPK activity in other experimental models can be reached in the CNS of SMA mice.

Next, we investigated the effects of MW150 on the phenotype of SMA mice, which fail to gain weight or right themselves and survive for an average of 14 days postnatally (49). MW150 was administered to SMA mice beginning at P0 via daily IP injections at a dose of 5mg/kg, which is the most effective dose established in previous mouse studies (46–48, 51–54). Importantly, treatment with MW150 improved motor function of SMA mice as measured by the righting time assay relative to vehicle treated SMA mice (Figure 4A). The effects on motor behavior were marked but deteriorated together with the overall health of SMA mice prior to death. Accordingly, MW150 treatment had no effect on weight gain or survival of SMA mice (Figure 4B and 4C), consistent with MW150 likely improving some but not all of the deficits underlying SMA pathology. We then tested whether MW150 treatment had any effect on expression of SMN in SMA mice. Western blot analysis did not reveal any effect of MW150 on SMN protein levels in the spinal cord of SMA mice (Figure 4D and 4E). Thus, MW150-mediated inhibition of p38 MAPK improves motor function of SMA mice without inducing SMN.

**Figure 4.**
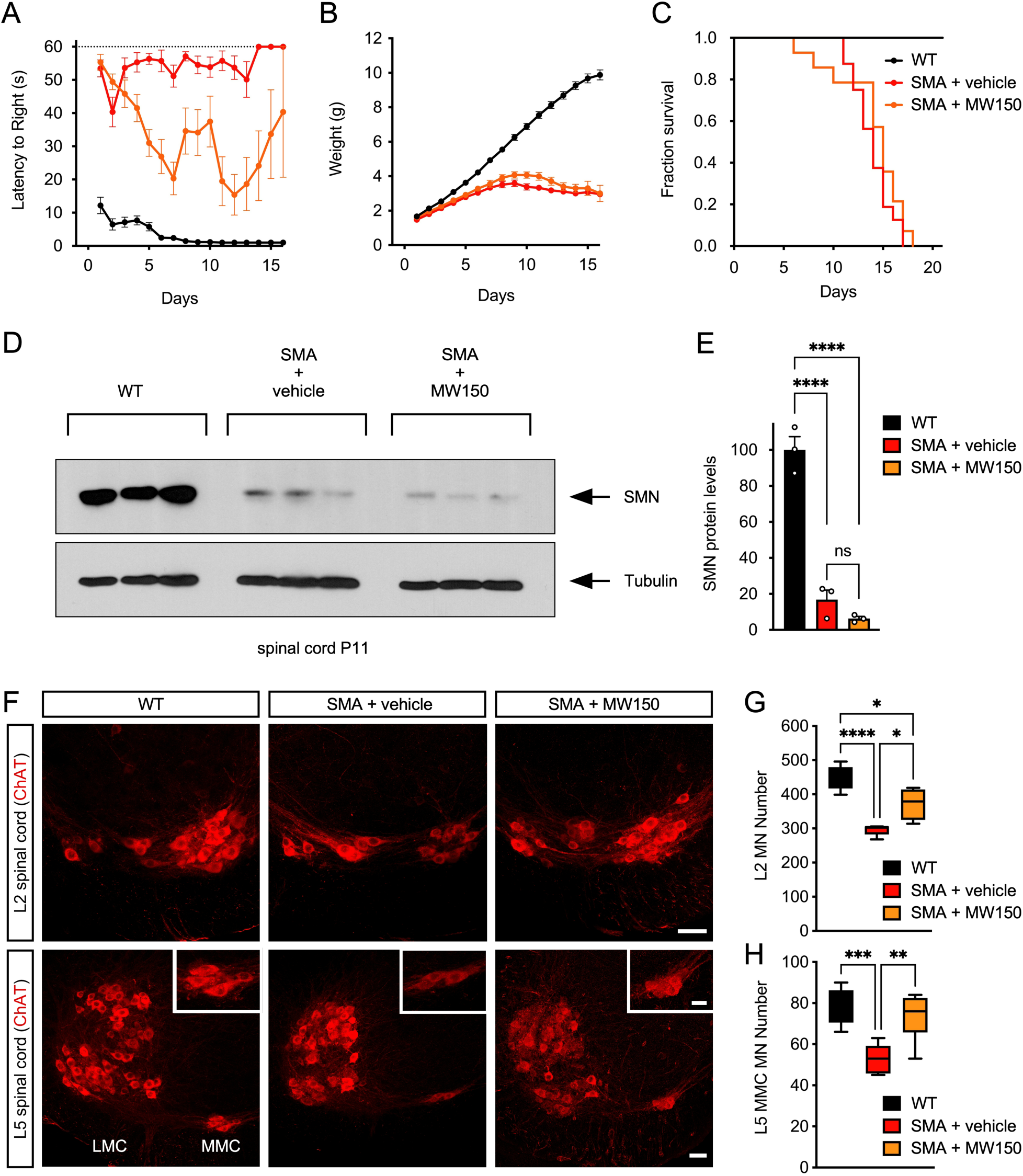
Pharmacological inhibition of p38 MAPK with MW150 improves motor behavior through neuroprotective effects in SMA mice. (**A-C**) Analysis of righting time (A), weight gain (B), and survival (C) of untreated WT mice (n=17) and SMA mice treated daily with vehicle (n=16) or 5mg/kg MW150 (n=14) starting at P0. (**D**) Western blot analysis of SMN levels in P11 spinal cords from the same experimental groups as in (A-C). Tubulin was used as loading control. (**E**) Quantification of SMN levels from the experiment in (D). Normalized mean, SEM, and individual values from 3 independent biological replicates are shown. One-way ANOVA and Tukey’s *post hoc* test. ****P < 0.0001; ns, not significant. (**F**) ChAT immunostaining of L2 and L5 spinal cords isolated at P11 from uninjected WT mice and SMA mice treated daily with vehicle or MW150 (5mg/kg) starting at P0. L5 lateral motor column (LMC) and medial motor column (MMC) motor neuron pools are indicated, and magnified views of L5 MMC motor neurons are shown in the insets. Scale bars = 50µm and 25µm (insets). (**G**) Total number of L2 motor neurons in the same experimental groups as in (F). The box and whiskers graph shows the median, interquartile range, minimum and maximum values from the following number of mice: WT (n=5), SMA+vehicle (n=5), and SMA+MW150 (P0) (n=4). One-way ANOVA with Tukey’s *post hoc* test. ****P < 0.0001; *P < 0.05. (**H**) Total number of L5 MMC motor neurons in the same experimental groups as in (F). The box and whiskers graph shows the median, interquartile range, minimum and maximum values from the following number of mice: WT (n=6), SMA+vehicle (n=6), and SMA+MW150 (P0) (n=6). One-way ANOVA with Tukey’s *post hoc* test. ***P < 0.001; **P < 0.01.

We previously reported that death of vulnerable motor neurons in SMA mice is driven by the convergence of stabilization and amino-terminal phosphorylation of p53 through distinct mechanisms (12, 13, 18), the latter of which is mediated by p38 MAPK activation and inhibited by MW150 (13). Therefore, we quantified the total number of lumbar L2 and L5 medial motor column (L5 MMC) SMA motor neurons, which are known to degenerate in the SMNβ7 model (12, 18, 19). Immunohistochemistry with anti-ChAT antibodies revealed the expected loss of both L2 and L5 MMC motor neurons in the lumbar spinal cord of vehicle treated SMA mice relative to WT controls at P11 (Figure 4F-H), while daily treatment with 5mg/kg of MW150 starting at P0 significantly rescued the survival of these vulnerable motor neuron pools (Figure 4F-H). Together with our previous study (13), these results support the conclusion that the phenotypic benefit of p38 MAPK inhibition on motor function is driven by neuroprotection of SMA motor neurons.

To address this point further, we analyzed the phenotypic effects of daily treatment with MW150 in Smn^2B/-^ SMA mice (55), which are characterized by severe synaptic pathology but, unlike SMNΔ7 SMA mice, do not display significant motor neuron death (56, 57). We found that MW150 had no effects on motor function in this mouse model of SMA (Supplementary Figure 2). This is consistent with pharmacological inhibition of p38 MAPK with MW150 acting selectively on the death pathway of SMA motor neurons.

### An experimental paradigm of combinatorial treatment for evaluation of neuroprotective effects in SMA mice

We next sought to investigate potential synergies of the neuroprotection afforded by MW150 treatment in combination with SMN upregulation. Pre-clinical studies have shown that SMN-inducing therapies are highly effective in rescuing the disease phenotype in mouse models and do not display the variability in clinical response that is found in the real world setting of SMA patients. Therefore, we set out to develop an experimental paradigm for suboptimal treatment with SMN inducing drugs in which to test the neuroprotective effects of p38 MAPK inhibition – an approach often used for combinatorial studies in SMA mice (58–64). Specifically, we tested the phenotypic response of SMA mice subjected to early or delayed SMN induction by treatment with SMN-C3. The paradigm of delayed treatment was preferred to that of lower doses of SMN-C3 starting from birth to clearly distinguish early, MW150-dependent effects without the confounding element of a concomitant moderate SMN induction.

We performed daily IP injections of SMN-C3 in SMA mice starting at either P0 (early treatment) or P8 (delayed treatment) using the previously established therapeutic dose of 3mg/kg. Consistent with previous studies (43), SMN-C3 early treatment induced a progressive, strong rescue of motor function assessed by the righting time (Figure 5A), a marked increase in weight gain (Figure 5B), and extension of survival relative to vehicle-treated SMA mice up to P30 (Figure 5C), which was set as the time of study termination. In contrast, SMA mice receiving delayed SMN-C3 treatment showed a modest yet variable improvement of motor function towards disease end stage (Figure 5A), but no significant increase in either weight gain or survival relative to vehicle-treated SMA mice (Figure 5B and 5C).

**Figure 5.**
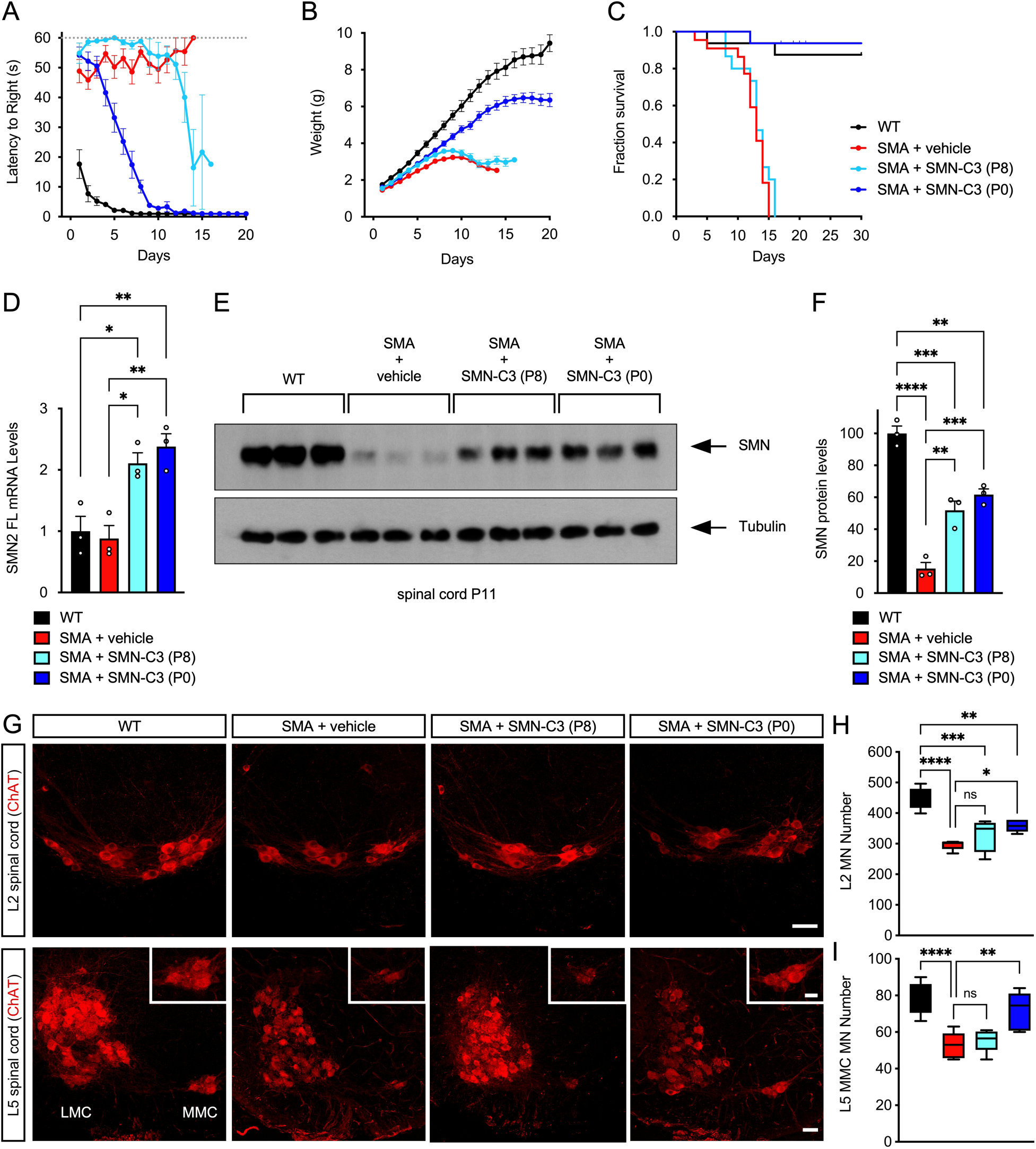
Analysis of delayed treatment of SMA mice with SMN-C3. **(A-C)** Analysis of righting time (A), weight gain (B), and survival (C) of untreated WT mice (n=16) and SMA mice treated daily with vehicle (n=22) or SMN-C3 (3mg/kg) starting at P0 (n=16) and P8 (n=15). Data represent mean and SEM. (**D**) RT-qPCR analysis of the levels of full-length *SMN2* mRNA in P11 spinal cords from the same experimental groups as in (A-C). Mean, SEM, and individual values normalized to WT samples as a control are shown (n=3 mice). One-way ANOVA followed by Tukey’s *post hoc* test. **P < 0.01; *P < 0.05. (**E**) Western blot analysis of SMN levels in P11 spinal cords from the same experimental groups as in (A-C). Tubulin was used as loading control. (**F**) Quantification of SMN levels from the experiment in (E). Normalized mean, SEM, and individual values from 3 independent biological replicates are shown. One-way ANOVA followed by Tukey’s *post hoc* test. ****P < 0.0001; ***P < 0.001; **P < 0.01. (**G**) ChAT immunostaining of L2 and L5 spinal cords isolated at P11 from uninjected WT mice and SMA mice treated daily with vehicle or SMN-C3 (3mg/kg) starting at P0 and P8. L5 LMC and MMC motor neuron pools are indicated, and magnified views of L5 MMC motor neurons are shown in the insets. Scale bars = 50µm and 25µm (insets). (**H**) Total number of L2 motor neurons in the same experimental groups as in (G). The box and whiskers graph shows the median, interquartile range, minimum and maximum values from the following number of mice: WT (n=5), SMA+vehicle (n=5), SMA+SMN-C3 (P0) (n=5), and SMA+SMN-C3 (P8) (n=4). One-way ANOVA with Tukey’s *post hoc* test. ****P < 0.0001; ***P < 0.001; **P < 0.01; *P < 0.05; ns, not significant. (**I**) Total number of L5 MMC motor neurons in the same experimental groups as in (G). The box and whiskers graph shows the median, interquartile range, minimum and maximum values from the following number of mice: WT (n=6), SMA+vehicle (n=6), SMA+SMN-C3 (P0) (n=6), and SMA+SMN-C3 (P8) (n=6). One-way ANOVA with Tukey’s *post hoc* test. ****P < 0.0001; **P < 0.01; ns, not significant.

To determine whether SMN-C3 induced comparable increases in SMN expression irrespective of the time of intervention, we analyzed the levels of both exon 7 inclusion in SMN2 mRNAs and SMN protein in the spinal cord of SMA mice at P11. RT-qPCR analysis showed a similar 2-fold increase in the spinal cord levels of full-length SMN2 mRNA following either early or delayed delivery of SMN-C3 in SMA mice relative to controls (Figure 5D). Moreover, SMN protein levels reached about 60% of normal in treated SMA mice regardless of the time SMN-C3 treatment was initiated (Figure 5E and F). These results indicate that SMN-C3 drug delivery is similarly effective in inducing SMN upregulation irrespective of the time of treatment initiation, and longer treatment does not result in greater SMN induction.

It is conceivable that the observed differences in phenotypic correction reflect accumulation of early damage that cannot be effectively repaired with delayed treatment (Figure 5A-C), making loss of motor neurons the most likely candidate because it is an irreversible pathogenic event after it has occurred. To address this possibility, we determined the effects of early and delayed treatment with SMN-C3 on the survival of lumbar L2 and L5 MMC motor neurons in SMA mice. Importantly, we found that SMN-C3 treatment at P8 did not prevent motor neuron loss (Figure 5G-I), which was instead significantly corrected by early treatment. Moreover, similar to MW150 (Figure 4F-H), early treatment with SMN-C3 was more effective in promoting the survival of L5 MMC than L2 SMA motor neurons (Figure 5G-I), which likely reflects differences in the timing of irreversible commitment to death of these distinct motor neuron pools (18, 19, 65).

Together, these results indicate that the lack of neuroprotection correlates with limited phenotypic improvement by delayed SMN induction in SMA mice and that our experimental paradigm of delayed intervention is suitable for evaluating neuroprotective drugs in combination with SMN-inducing therapies.

### Synergistic improvement of the SMA phenotype by MW150 in combination with delayed SMN upregulation

To evaluate whether MW150-dependent neuroprotection provides additional phenotypic benefit relative to delayed SMN induction in SMA mice, we analyzed the phenotypic response of SMA mice that were treated daily with vehicle or MW150 (5mg/kg) starting at P0 in combination with SMN-C3 (3mg/kg) treatment at P8 (Figure 6A). Under these conditions, we found remarkable added benefit of combinatorial treatment with MW150 relative to vehicle (Figure 6B-D). The improvement in motor function was much more rapid and robust in MW150-treated animals relative to SMA mice treated with delayed SMN-C3 alone (Figure 6B). We also observed a strong increase in weight gain (Figure 6C), which either treatment alone fails to induce (compare with Figures 4B and 5B), and a notable extension of survival in about half of the combinatorially treated SMA mice (Figure 6D). Thus, MW150 treatment strongly enhances the phenotypic benefit of delayed SMN upregulation in SMA mice, highlighting the key impact that preserving motor neuron survival may exert on the potential for correction of SMA pathology by SMN inducing drugs.

**Figure 6.**
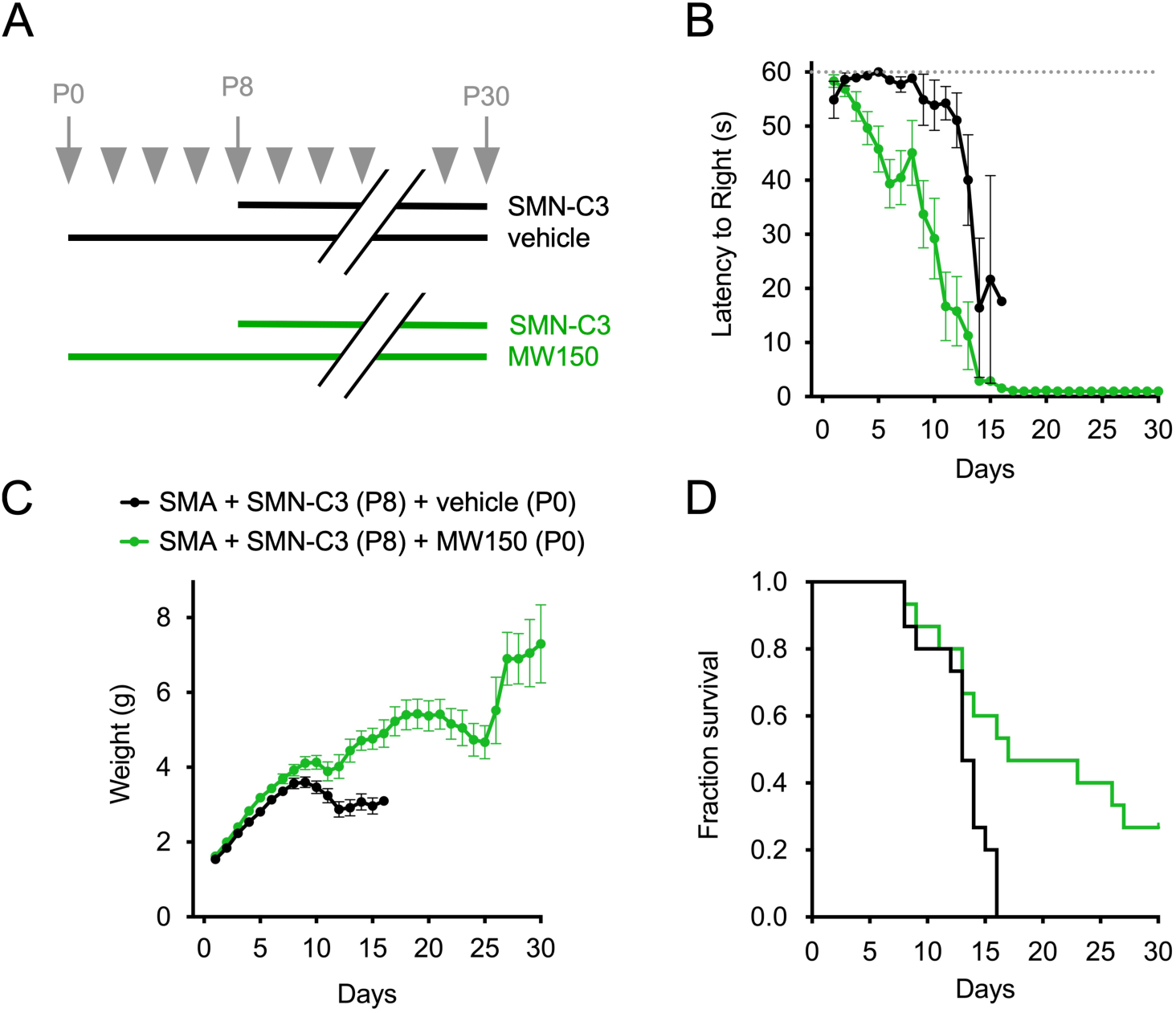
Phenotypic analysis of combinatorial treatment of SMA mice with MW150 and SMN-C3. (**A**) Schematic of the experimental design for delayed SMN-C3 treatment alone and combinatorial treatment with both SMN-C3 and MW150. SMA mice received either daily treatment with vehicle (delayed SMN-C3) or 5mg/kg MW150 starting at P0 (combinatorial treatment). Daily administration of 3mg/kg SMN-C3 initiated at P8. (**B-D**) Analysis of righting time (B), weight gain (C), and survival (D) of SMA mice treated daily with delayed SMN-C3 (n=15) or the combinatorial treatment (n=15). Data in B and C represent mean and SEM. Kaplan-Meyer analysis of survival is shown in (D).

### Neuroprotection by MW150 allows synaptic rewiring of motor neurons by delayed SMN induction

We next sought to establish the basis for the observed synergistic, beneficial effects of MW150 treatment in combination with delayed SMN-C3 treatment. We and others have recently shown that motor neuron deafferentation and neuromuscular junction (NMJ) denervation are shared pathogenic features across all SMA mouse models, while motor neuron death is a selective signature of SMNΛ17 SMA mice (56, 57). We reasoned that, while delayed treatment with SMN-C3 is ineffective in rescuing vulnerable motor neurons from early commitment to the death pathway (Figure 5G-I), prior treatment with MW150 rescues these motor neurons (Figure 4F-H) and could provide the opportunity for SMN-C3 to functionally correct other, p38 MAPK-independent pathological features of SMA. Specifically, we investigated the possibility of synaptic rewiring of motor neurons both centrally - with other neurons of the motor circuit such as VGluT1^+^ proprioceptors - and distally - through NMJ reinnervation of skeletal muscle.

We first analyzed neuromuscular junction (NMJ) innervation in the vulnerable, axial muscle *quadratus lumborum* (QL) of SMA mice at P11 by immunohistochemistry and confocal microscopy. Pre-synaptic terminals of motor neurons were stained with antibodies against Neurofilament-M and Synaptophysin, while post-synaptic motor endplates were stained with fluorescently labeled bungarotoxin (Figure 7A). Consistent with previous studies (12, 14, 18), we found that ∼30% of the NMJs are fully denervated in SMA mice at P11 (Figure 7A and 7B). Importantly, while this defect was not corrected by treatment with either MW150 alone or delayed SMN-C3, the combinatorial therapy significantly improved NMJ innervation (Figure 7A and B), albeit not as effectively as early SMN-C3 treatment. Thus, while NMJ denervation in SMA mice is independent from p38 MAPK activation, the neuroprotective effects of MW150 enables early improvement of synaptic connections between motor neuron and muscle by delayed SMN induction.

**Figure 7.**
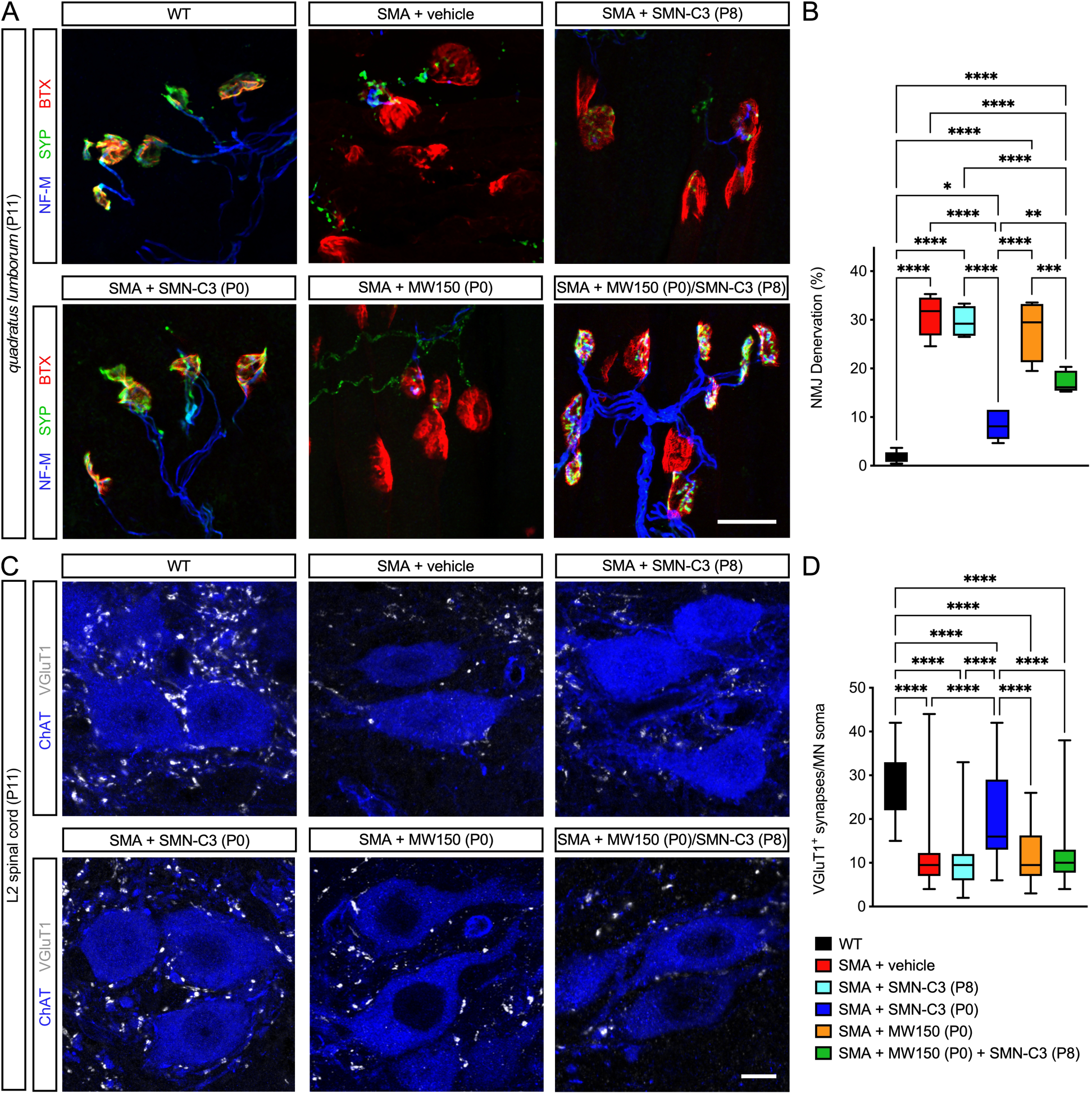
MW150 treatment promotes partial improvement of NMJs but not central synapses by delayed SMN induction at early times. (**A**) NMJ staining with bungarotoxin (BTX, red), synaptophysin (Syp, green), and neurofilament (NF-M, blue) of QL muscles isolated at P11 from uninjected WT mice and SMA mice treated with vehicle, MW150, early (P0) and delayed (P8) SMN-C3 either alone or in combination with MW150. Scale bar = 25 µm. (**B**) Percentage of fully denervated NMJs in the QL muscle from the same experimental groups as in (A). The box and whiskers graph shows the median, interquartile range, minimum and maximum values from the following number of mice: WT (n=8), SMA+vehicle (n=8), SMA+MW150 (P0) (n=4), SMA+SMN-C3 (P0) (n=6), SMA+SMN-C3 (P8) (n=5), and SMA+SMN-C3 (P8)/MW150 (P0) (n=5). (**C**) Immunostaining of VGluT1^+^ synapses (gray) and ChAT^+^ motor neurons (blue) of L2 spinal cords isolated at P11 from the same experimental groups as in (A). Scale bar = 10 µm. (**D**) Number of VGluT1^+^ synapses on the somata of L2 motor neurons from the same experimental groups as in (A). The box and whiskers graph shows the median, interquartile range, minimum and maximum from the following number of motor neurons (n) from 4 mice for each experimental group: WT (n=39), SMA+vehicle (n=50), SMA+MW150 (P0) (n=46), SMA+SMN-C3 (P0) (n=47), SMA+SMN-C3 (P8) (n=52), and SMA+SMN-C3 (P8)/MW150 (P0) (n=42). Statistics in (B) and (D) were performed with one-way ANOVA with Tukey’s *post hoc* test. ****P < 0.0001; ***P < 0.001; **P < 0.01; *P < 0.05.

We then investigated the effects of combinatorial treatment on motor neuron deafferentation. Analysis by immunohistochemistry with ChAT and VGluT1 antibodies followed by confocal microscopy showed that the number of VGluT1^+^ proprioceptive synapses impinging on the somata of lumbar motor neurons are strongly reduced in SMA mice relative to normal controls at P11 (Figure 7C and 7D), consistent with previous studies (13, 19, 20, 66). Interestingly, neither delayed treatment of SMA mice with SMN-C3 nor MW150 treatment either alone or in combination increased the number of proprioceptive synapses (Figure 7C and 7D). In contrast, early SMN-C3 treatment significantly improved motor neuron deafferentation (Figure 7C and 7D). These results indicated that the loss of VGluT1^+^ synapses in SMA mice is independent from p38 MAPK activation and that sensory-motor synaptic connectivity cannot be rapidly reestablished by delayed SMN induction.

To determine whether prolonged SMN induction could further improve synaptic rewiring of SMA motor neurons, we analyzed proprioceptive synapses and NMJ innervation at P21 in SMA mice treated with the combinatorial therapy. WT and early SMN-C3 treated SMA mice were used as controls, while SMA mice treated with vehicle, MW150 alone, and delayed SMN-C3 could not be investigated because they die prior to this time point (Figures 4C and 5C). Remarkably, we found no significant differences in the degree of muscle innervation across experimental groups with essentially all NMJs of the vulnerable QL being fully innervated (Figure 8A and 8B). Moreover, the number of VGluT1^+^ synapses on motor neurons from SMA mice treated with the combinatorial therapy was similar to that of mice treated early with SMN-C3 and not significantly different from WT mice (Figure 8C and 8D). Thus, extending SMN-C3 treatment from P11 to P21 not only promoted full NMJ innervation but also the rewiring of proprioceptive synapses on motor neurons, counteracting their early loss (compare Figures 7D and 8D).

**Figure 8.**
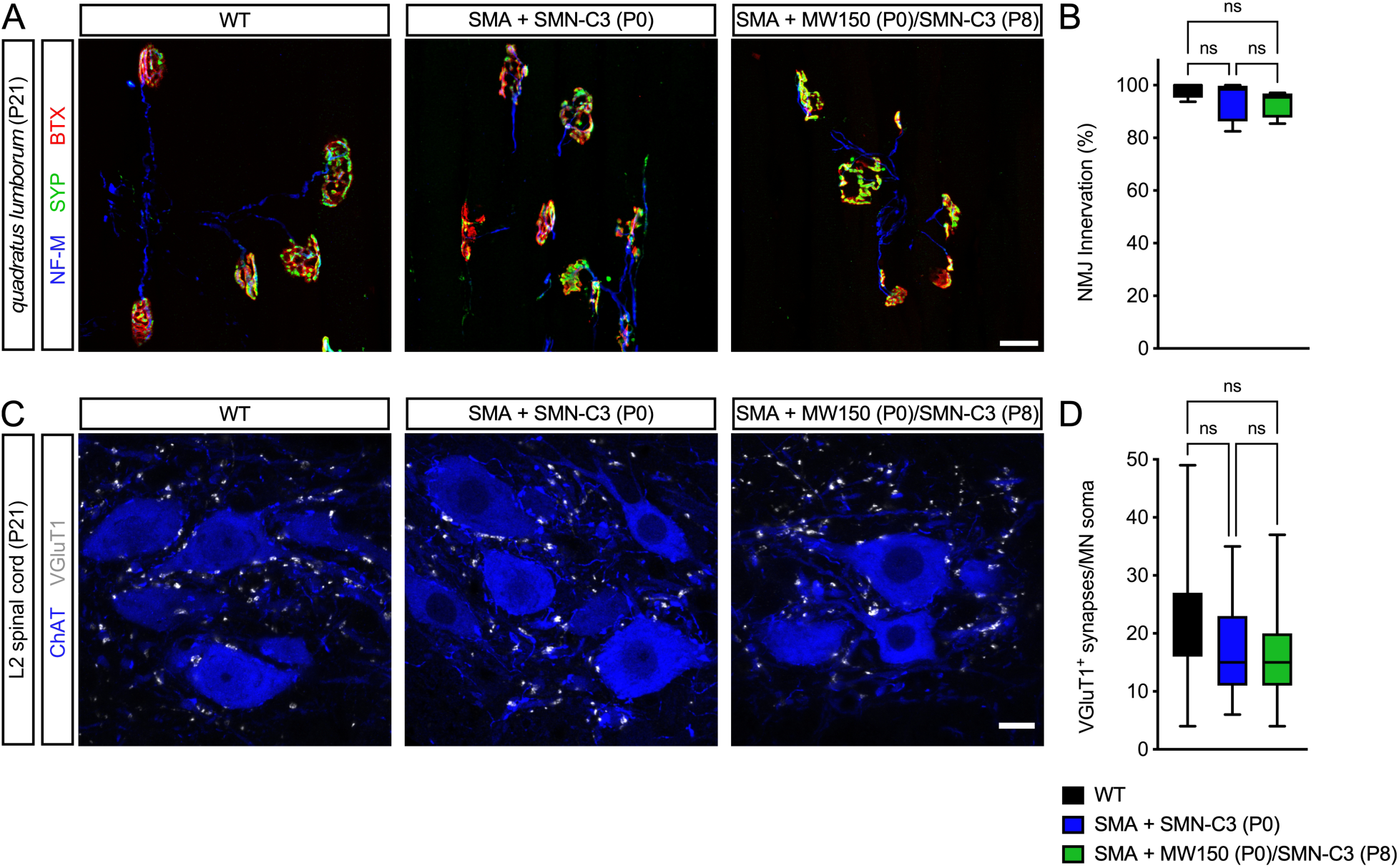
MW150 treatment promotes restoration of NMJs and central synapses by delayed SMN induction at later times. (**A**) NMJ staining with bungarotoxin (BTX, red), synaptophysin (Syp, green), and neurofilament (NF-M, blue) of QL muscles isolated at P21 from uninjected WT mice, SMA mice injected daily with SMN-C3 (3mg/kg) starting at P0, and SMA mice injected with SMN-C3 (3mg/kg) at P8 in combination with MW150 (5mg/kg) starting at P0. Scale bar = 25 µm. (**B**) Percentage of fully innervated NMJs in the QL muscle from the same experimental groups as in (A). The box and whiskers graph shows the median, interquartile range, minimum and maximum values from the following number of mice: WT (n=4), SMA+SMN-C3 (P0) (n=4), and SMA+SMN-C3 (P8)/MW150 (P0) (n=4). (**C**) Immunostaining of VGluT1^+^ synapses (gray) and ChAT^+^ motor neurons (blue) of L2 spinal cords isolated at P21 from the same experimental groups as in (A). Scale bar = 25 µm. (**D**) Number of VGluT1^+^ synapses on the somata of L2 motor neurons from the same experimental groups as in (A). The box and whiskers graph shows the median, interquartile range, minimum and maximum values from the following number of neurons (n) from 4 mice for each experimental group: WT (n=38), SMA+SMN-C3 (P0) (n=38), and SMA+SMN-C3 (P8)/MW150 (P0) (n=39). Statistics in (B) and (D) were performed with one-way ANOVA with Tukey’s *post hoc* test. ns, not significant.

Collectively, these results demonstrate that the neuroprotective effects of MW150 can synergize with SMN inducing drugs and provide the opportunity to enhance the therapeutic impact of delayed SMN upregulation by promoting synaptic rewiring of SMA motor neurons within spinal sensory-motor circuits. Provided that motor neurons are spared from death, these findings show that even delayed SMN induction can restore functional synaptic connections with proprioceptive neurons and skeletal muscle after their loss, indicating that they are reversible pathogenic events with distinct kinetics for SMN-dependent correction.

## Discussion

The identification of cellular factors and pathways that contribute to pathogenic events downstream of SMN deficiency is key to understand disease mechanisms and broaden the range of targets for developing SMA therapies that can complement SMN upregulation. Here, we performed a phenotypic screen for chemical modifiers of SMN biology and identified inhibitors of p38 MAPK as suppressors of cell proliferation defects induced by SMN deficiency in mouse fibroblasts. We further show that SMN deficiency induces p38 MAPK activation both *in vitro* and *in vivo,* and that pharmacological inhibition of this pathway improves motor function in SMA mice through SMN-independent neuroprotective effects. Using a highly optimized p38αMAPK inhibitor (MW150) and a specific paradigm of combinatorial treatment, we demonstrate synergistic enhancement of the phenotypic benefit induced in SMA mice by MW150 together with an SMN inducing drug relative to either treatment alone. By promoting survival of SMA motor neurons, pharmacological inhibition of p38 MAPK synergizes with SMN induction and enables enhanced synaptic rewiring within sensory-motor circuits, which results in increased motor function, weight gain, and survival. Together, our studies identify the p38 MAPK pathway as a therapeutic target and MW150 treatment as a candidate pharmacological approach for SMN-independent neuroprotection with the potential of enhancing the clinical benefit of SMN induction in SMA.

Unbiased phenotypic screens have the power to identify disease modifiers but have not yet been employed as discovery tools to identify disease mechanism-relevant therapeutic targets in SMA. To fill this gap, we performed a cell-based phenotypic screen designed to capture molecules that promote SMN expression or act downstream of SMN depletion. Interestingly, none of the identified hits increase SMN expression, but may function through mechanisms that are SMN independent and, in principle, suitable for use in combination with approved SMA therapies. We prioritized our follow up studies focusing on p38 MAPK because i) EO1428 was the top validated hit of our screen, ii) chemically different p38 MAPK inhibitors similarly promoted proliferation of SMN-deficient mouse fibroblasts, and iii) a highly selective, optimized compound (MW150) was available for *in vivo* studies (46–48, 51–54). Furthermore, the p38 MAPK family comprises a group of four serine-threonine protein kinases (α, β, ο, and ψ) that process a variety of stimuli and stress signals through phosphorylation cascades and are involved in cell proliferation, differentiation, and death (67). Thus, the effects of p38 MAPK inhibitors in promoting cell proliferation of SMN-deficient cells are consistent with the biological role of this pathway. Importantly, accumulation of phosphorylated p38 MAPK in both NIH3T3 cells and the spinal cord of SMA mice provides direct evidence of its activation by SMN deficiency.

To demonstrate the relevance of p38 MAPK in SMA pathology, we investigated the effects of its pharmacological inhibition using MW150. Our findings confirm the outstanding brain permeability of MW150 and show that treatment with this small molecule improves motor function of SMA mice in an SMN-independent manner. Mechanistically, p38 MAPK inhibition has neuroprotective effects by acting on the degenerative pathway of SMA motor neurons. We have previously implicated upregulation and phosphorylation of the amino-terminal transcriptional activation domain of the tumor suppressor p53 as two distinct pathogenic events that converge to drive the death of vulnerable motor neuron pools in SMA mice (18). We have also shown that both events originate from disruption of SMN’s function in snRNP assembly and downstream dysregulation of specific splicing events (12, 13). On one hand, alternative splicing changes of specific exons in Mdm2 and Mdm4 pre-mRNAs drive p53 stabilization (12). On the other, dysregulated U12 splicing of Stasimon promotes phosphorylation of p53 (13). Importantly, the latter involves p38 MAPK activation whose inhibition by MW150 reduces both p53 phosphorylation and motor neuron death in SMA mice.

Building on the neuroprotective effects of MW150, we explored the therapeutic potential of pharmacological inhibition of the p38 MAPK pathway as a candidate disease-modifying approach for use in combination with SMN-inducing drugs. To do so, we employed an experimental paradigm for suboptimal treatment of SMA mice in which delayed SMN upregulation neither improves motor neuron survival nor synaptic pathology, resulting in the lack of significant phenotypic rescue. Importantly, we document strong synergistic effects when MW150 treatment is used in combination with delayed SMN induction, leading to an improvement of the SMA phenotype that far exceeds that of each treatment alone. The synergy stems from the compounded effects of i) MW150 promoting motor neuron survival, and ii) SMN-C3 improving other p38 MAPK-independent SMA deficits. These findings provide direct experimental support to the importance of sparing motor neurons from death in order to achieve the most effective correction of SMA pathology by SMN inducing drugs. In addition to being a disease hallmark (1, 2), motor neuron loss is an irreversible pathogenic event that cannot be corrected after it has occurred and one that, if insufficiently addressed, can limit the clinical benefit of current SMN inducing therapies (29, 30, 33). In this context, the synergistic effects elicited by pharmacological inhibition of p38 MAPK together with SMN inducing drugs have clinical implications for combinatorial SMA therapy.

Our study provides proof-of-concept that preserving motor neuron survival enables enhanced synaptic rewiring of SMA motor neurons within spinal sensory-motor circuits and increases phenotypic benefit by SMN upregulation. It also yields insights into the potential for therapeutic correction of synaptic pathology by delayed SMN induction. Although increased NMJ innervation and proprioceptive synaptic inputs on motor neurons have been documented in SMA mice following SMN restoration (13, 58, 68–71), these previous studies with early initiation of therapeutic intervention could not clearly distinguish whether SMN induction can act by preventing or slowing down synaptic loss versus reestablishing lost connections. Using our experimental paradigm for combination treatment, we show that delayed SMN induction can promote the formation of new synaptic connections between motor neurons and either proprioceptive neurons or skeletal muscle after they are lost due to SMA pathology. Thus, NMJ denervation and proprioceptive deafferentation are reversible pathogenic events that, unlike motor neuron death, can be recovered even at late symptomatic stages of the disease. Of note, we also show that proprioceptive synapses on motor neurons are reestablished significantly more slowly than NMJs in SMA mice, revealing that different synaptic connections follow distinct kinetics of restoration after initiation of treatment with SMN inducing drugs. Collectively, these findings highlight the mechanisms by which the neuroprotective effects of p38 MAPK inhibition can synergize with SMN upregulation to enhance therapeutic efficacy in SMA.

The p38 MAPK pathway has previously been considered a potential therapeutic target for inflammatory diseases and several neurological disorders (72, 73). Accordingly, p38 MAPK inhibitors have been tested in cellular and animal models of late onset neurodegeneration (46, 47, 52, 74–80). Until recently, however, validation of p38 MAPK as a drug discovery target has been limited by the lack of highly selective inhibitors amenable to *in vivo* use in the CNS (81). The development of MW150 has changed this scenario. MW150 is a potent and highly selective inhibitor of p38α MAPK that has previously been shown to be effective in mouse models of AD and other CNS disorders (46–48, 51–54). MW150 is a water-soluble and orally bioavailable p38αMAPK inhibitor that has been optimized through medicinal chemistry and exhibits excellent CNS permeability, metabolic stability, safety, and reduced risk for drug-drug interaction. Thus, MW150 has all the desired drug-like properties for the treatment of neurological disorders and is currently under clinical testing. Together with our findings, the favorable pharmacological properties of MW150 should facilitate its entry into the clinical pipeline for testing as a potential combination treatment with SMN inducing drugs for SMA.

In conclusion, our studies support the therapeutic potential of pharmacologically targeting the p38 MAPK pathway as a candidate, SMN-independent neuroprotective approach to enhance motor neuron survival and the clinical benefit of SMN-inducing therapies in SMA.

## Materials and Methods

### Cell lines

The NIH3T3 cell lines used in this study were previously described. NIH3T3-Smn_RNAi_ cells express the tetracycline-dependent repressor (TetR) regulating the expression of an shRNA targeting endogenous mouse Smn driven by an H1_TO_ promoter harboring two tandem copies of the Tet operator sequence (10, 11). NIH3T3-SMN2/Smn_RNAi_ cells were derived from NIH3T3-Smn_RNAi_ to express low levels of RNAi-resistant human SMN from a stably integrated human *SMN2* gene (42). NIH3T3-SmB_RNAi_ cells express TetR and an shRNA targeting endogenous mouse SmB driven by the H1_TO_ promoter and have normal SMN levels (10). All NIH3T3 cell lines were cultured in Dulbecco’s Modified Eagle’s Medium (DMEM) with high glucose (Gibco) containing 10% fetal bovine serum (HyClone), 2mM L-glutamine (Gibco), and 0.1mg/ml gentamycine (GIBCO). RNAi was induced by addition to the growth medium of Doxycycline HCl (Fisher Scientific) at the final concentration of 100 ng/ml.

### Cell proliferation assay

The cell proliferation assay in 96-well format was performed as previously described with minor modifications (42). NIH3T3 cells were cultured with Dox for 5 days prior to seeding in 96-well optical plates (Greiner Bio-One) at 100 cells per well in 200μl of Dox-containing media. Compounds were added 4 hours post-plating and experiments terminated after 5 additional days in culture by addition of a 4X solution containing 16% paraformaldehyde and 8mg/ml of Hoechst in PBS to each well of the 96-well plate and further incubation for 15 minutes at room temperature. After washing with PBS, cell number determination was performed by whole-well imaging acquisition using a plate imaging system (TROPHOS Plate RUNNER HD) followed by nuclear counting with the TINA software (TROPHOS).

### Chemical compounds

For the chemical screen, we used a library of 1,307 small molecules containing the complete Tocriscreen Mini (Tocris, #2890) collection and additional compounds from Tocris highly enriched in FDA-approved drugs, known signaling inhibitors, and other bioactive molecules. SMN-C3 was custom synthesized by Combi-Block, Inc. according to the published compound structure (43), dissolved at a concentration of 1.2mg/ml in 100% DMSO, and stored in aliquots at −80°C until use. MW150 was synthesized as previously described (46), dissolved at a concentration of 10mg/ml in sterile 0.9% saline, and stored in aliquots at room temperature until use. All other compounds were obtained from Tocris Bioscience and dissolved in 100% DMSO unless otherwise specified.

### Cell-based chemical screen

A phenotypic screen for chemical compounds that promote cell proliferation of SMN-deficient mouse fibroblasts was carried out using Dox-treated NIH3T3-SMN2/Smn_RNAi_ cells. A Tecan Freedom EVO 200 robotic platform equipped with two-arm operation system and 96-channel pipetting stations was employed for liquid handling. Several parameters of the cell-based assay including cell density, the time of Dox pre-treatment prior to plating and from compound addition to readout, inter-well and inter-plate variability, and edge effects were optimized prior to screening implementation. For the chemical screen, NIH3T3-SMN2/Smn_RNAi_ cells were cultured with Dox for 5 days prior to seeding in 96-well optical plates (Greiner Bio-One) at 100 cells per well in 200μl of Dox-containing media. 1,307 compounds from the Tocris library were tested at a single concentration (10 μM) and one compound per well was added 4 hours post-plating. In each 96-well plate, 6 wells were reserved for negative drug controls (0.5% DMSO) and 6 wells for positive drug controls (500nM SMN-C3). Following 5 days of incubation, each well was scored using Hoechst staining and direct determination of cell number by imaging with the TROPHOS system. Any plate in which all of positive control values did not exceed the range of negative control values were rejected. The primary screen was carried out in triplicate plates and hits were defined as compounds causing a mean increase in cell number greater than the mean cell number of vehicle-treated cells plus 3 standard deviations. The screen yielded 15 compounds as initial hits, which underwent additional validation by i) visual inspection of 96-well images to check for drug precipitates that may affect readout, ii) repeat testing in multiple replicates, iii) dose-response analysis, and iv) counter-screen assays. Primary hits were initially retested at 10 µM in 6 replicate wells using the same source of compounds and assay in NIH3T3-SMN2/Smn_RNAi_ cells. Confirmed compounds were then repurchased from the vendor and analyzed in dose-response studies using concentrations spanning those used in the initial screening and 6 replicates per concentration in NIH3T3-SMN2/Smn_RNAi_ cells. As counter-screen assays designed to reveal whether compounds promote cell proliferation in a non-specific manner, compounds were also tested for their effects on proliferation using Dox-treated NIH3T3-SmB_RNAi_ cells and NIH3T3-SMN2/Smn_RNAi_ cells cultured with either normal (10%) or reduced (2%) serum in the absence of Dox. The half maximal effective concentration (EC50) was estimated by non-linear curve fitting using the dose-response data and the GraphPad Prism 10 software. Validated active compounds were defined as those with a reproducible activity and a potency of EC50<10 µM.

### Mouse lines

All mouse work was performed in accordance with the National Institutes of Health Guidelines on the Care and Use of Animals, complied with all ethical regulations and was approved by the IACUC committee of Columbia University. Mice were housed in a 12h/12h light/dark cycle with access to food and water *ad libitum*. FVB.Cg-*Grm7^Tg(SMN2)89Ahmb^ Smn1^tm1Msd^* Tg(SMN2*delta7)4299Ahmb/J (JAX Strain #005025) mice were interbred to obtain SMA mutant mice (Le et al., 2005). The *Smn*^2B/2B^ mice on a pure FVB/N genetic background were previously described (55). *Smn^2B/2B^*mice were crossed with heterozygous *Smn^+/−^* mice harboring the *Smn1^tm1Msd^* knockout allele (82) on a pure FVB/N genetic background to generate *Smn^2B/−^* SMA mice and *Smn^2B/+^* control littermates as previously described (56). Genotyping was performed from tail DNA as previously described for the SMNΔ7 (8) and *Smn*^2B^ (56) mouse lines, respectively. Equal proportions of mice of both sexes were used and aggregated data are presented because gender-specific differences were not found.

### Drug treatments and behavioral analysis

SMA mice were treated daily via intraperitoneal (IP) injection with MW150 (5 mg/kg) starting at P0. SMN-C3 was administered IP daily at 3 mg/kg beginning at either P0 or P8 for early and delayed treatment, respectively. In the combinatorial treatment, SMA mice received 5 mg/kg MW150 starting at P0 and 3 mg/kg SMN-C3 from P8, with injections on alternating abdominal sides and at separate times. Mice from all experimental groups were monitored daily for survival and weight from birth for up to 30 days. Righting reflex was assessed by placing the mouse on its back and measuring the time it took to turn upright on its four paws (righting time). Cut-off test time was 60 seconds. For each testing session, the test was repeated three times, and the mean of the recorded times was calculated. The hindlimb suspension test was performed daily from P11 onward as previously described (56). Litters were culled to 6 pups and investigators performing behavioral assays were blinded to genotype and treatment.

### MW150 biodistribution in mice

The exposure levels of MW150 in plasma and brain of SMA mice was measured following a single IP injection of four escalating drug concentrations (2.5, 5, 10 and 20mg/kg) at P10. Brain and plasma samples were collected 3 hours after injection from isoflurane anesthetized mice, quickly frozen, and shipped to Absorption Systems for determination of MW150 levels by a previously developed LC-MS/MS analytical method (46).

### RNA analysis

RNA from whole spinal cord tissue was extracted with TRIzol reagent (Invitrogen) and treated with DNase I (Ambion). RNA was reverse transcribed using the RevertAid first-strand cDNA kit (Thermo Scientific). Quantitative PCR (qPCR) was carried out using Power SYBR green PCR master mix (Applied Biosystems) in QuantStudio 3 (Applied Biosystems). The expression levels of total (SMN2_TOT_) and full-length (SMN2_FL_) SMN mRNAs specifically transcribed from the *SMN2* gene were quantified as previously described (10). The following primers were used: SMN2_FL_ Fwd 5’-CACCACCTCCCATATGTCCAGATT-3’; SMN2_FL_ Rev 5’-GAATGTGAGCACCTTCCTTCTTT-3’; SMN2_TOT_ Fwd 5’-GTGAGGCGTATGTGGCAAAAT-3’; and SMN2_TOT_ Rev 5’-CATATAGAAGATAGAAAAAAACAGTACAATGACC-3’.

### Protein analysis

For Western blot analysis, mice were sacrificed and spinal cord collection was performed in a dissection chamber under continuous oxygenation (95%O_2_/5%CO_2_) in the presence of cold (∼12°C) artificial cerebrospinal fluid (aCSF) containing 128.35mM NaCl, 4mM KCl, 0.58mM NaH_2_PO_4_, 21mM NaHCO_3_, 30mM D-Glucose, 1.5mM CaCl_2_, and 1mM MgSO_4_. Total protein extracts were generated by homogenization of spinal cords in SDS sample buffer (2% SDS, 10% glycerol, 5% ß-mercaptoethanol, 60mM Tris-HCl pH 6.8, and bromophenol blue), followed by brief sonication and boiling. Proteins were quantified using the *RC DC*^TM^ Protein Assay (Bio-Rad) and analyzed by SDS/PAGE on 12% polyacrylamide gels followed by Western blotting as previously described (10). A list of the antibodies used is shown in Table S1.

### Immunohistochemistry

For morphological studies by immunohistochemistry, mice were deeply anesthetized using Avertin, the depth of anesthesia was checked by the toe pinch reflex, and transcardial perfusion was then performed with a saline solution followed by 4% paraformaldehyde (PFA). The spinal cord and skeletal muscles were dissected and post-fixed by immersion in 4% PFA overnight at 4°C. For immunohistochemistry, the spinal cords were briefly washed with PBS and the L2 and L5 lumbar segments were identified by the ventral roots, dissected, and subsequently embedded in warm 5% agar. Transverse sections (75μm) of the entire spinal segment were obtained with a VT1000 S vibratome (Leica). All the sections were then incubated overnight at room temperature with different combinations of primary antibodies diluted in PBS-T. The following day, six washing steps of 10 minutes each were done prior to incubation with secondary antibodies for 3 hours in PBS. Another six washing steps were performed before sections were mounted in 30% glycerol/PBS. For NMJ analysis, skeletal muscles were cryoprotected through sequential immersion in 10% and 20% sucrose/0.1M phosphate buffer (PB) for 1 hour at 4°C followed by overnight immersion in 30% sucrose/0.1M PB at 4°C. The following day, muscles were frozen embedded in Optimal Cutting Temperature (OCT) compound (Fisher), frozen on dry ice, and stored at −80°C until processing. Longitudinal cryosections (30μm) were collected onto Superfrost Plus glass slides (Fisher) using a CM3050S cryostat (Leica). Sections were washed once with PBS for 5 minutes to remove OCT, blocked for 1 hour with 5% donkey serum in TBS containing 0.2% Triton-X at room temperature and incubated with primary antibodies in blocking buffer overnight at 4°C. Following incubation, sections were washed three times for 10 minutes in TBS-T and then incubated with tetramethylrhodamine-conjugated α-bungarotoxin (Invitrogen #T1175, 1:500) and the appropriate secondary antibodies for 1 hour at room temperature, followed by 3 washing steps. Slides were mounted using Fluoromount-G Mounting Medium (SouthernBiotech). A list of the antibodies used is shown in Table S1.

### Confocal microscopy and image analysis

All images were collected with an SP5 confocal microscope (Leica) running the LAS AF software (v2.5.2.6939) and analyzed off-line using the Leica LAS X software (v1.9.0.13747). For motor neuron number quantification, 1024 x 1024 pixels images were acquired from all the 75μm sections of L2 and L5 spinal segments using a 20X objective at 3μm steps in the z-axis and a 200 Hz acquisition rate. Only motor neurons (ChAT^+^) with a clearly identifiable nucleus were counted to avoid double counting from adjoining sections. For quantification of VGluT1^+^ synapses, 1024 x 1024 pixels images were acquired from L2 spinal sections (75μm) using a 40X objective at 0.3μm steps in the z-axis and a 200 Hz acquisition rate. The total number of VGluT1^+^ synapses per soma was determined by counting all the corresponding inputs over the entire surface of each ChAT^+^ motor neuron cell body. At least 10 motor neurons per mouse were quantified. For NMJ analysis, 1024 x 1024 pixels images were acquired from 30μm muscle sections using a 20X objective at 2μm steps in the z-axis and a 200 Hz acquisition rate. Maximum intensity projections of confocal stacks were analyzed and at least 200 randomly selected NMJs per muscle were quantified. NMJs lacking any coverage of the α-bungarotoxin-labeled postsynaptic endplate by the presynaptic markers Synaptophysin and Neurofilament-M were scored as denervated and those with less than 50% coverage were scored as partially innervated.

### Statistical analysis

The sample size for each experiment is detailed in the figure legends. Statistical analysis was performed by two-tailed unpaired Student’s *t*-test or by two-way ANOVA followed by the Tukey’s multiple comparison test as indicated. Comparison of survival curves was performed using the Log-rank (Mantel-Cox) test. GraphPad Prism 10 for macOS Version 10.4.1 was used for all statistical analyses and P values are indicated as follows: *P<0.05; **P<0.01; ***P<0.001; **** P< 0.0001.

## Acknowledgements

We are grateful to George Mentis for helpful discussions and critical reading of the manuscript. We thank Rashmi Kothary for providing the *Smn^2B^* mouse line. This work was supported by grants from CureSMA (L.P.) and NS071092 (L.P.), NS083831 (L.P.), NS098363 (L.P.), NS102451 (L.P.) and NS116400 (L.P.) from NINDS.

## Data availability

The data presented in this study are available upon reasonable request from the corresponding author.

## Authors contributions

LP conceived and supervised the study; MJC, JED, MVA, ST, EW, IT, MKT and SY performed the experiments; MJC, JED, MVA, ST, EW, IT, MKT, SY and LP analyzed the data; CEH and DMW provided resources and experimental advice; MJC and LP wrote the paper with contributions from all authors.

---

**Figure S1.**
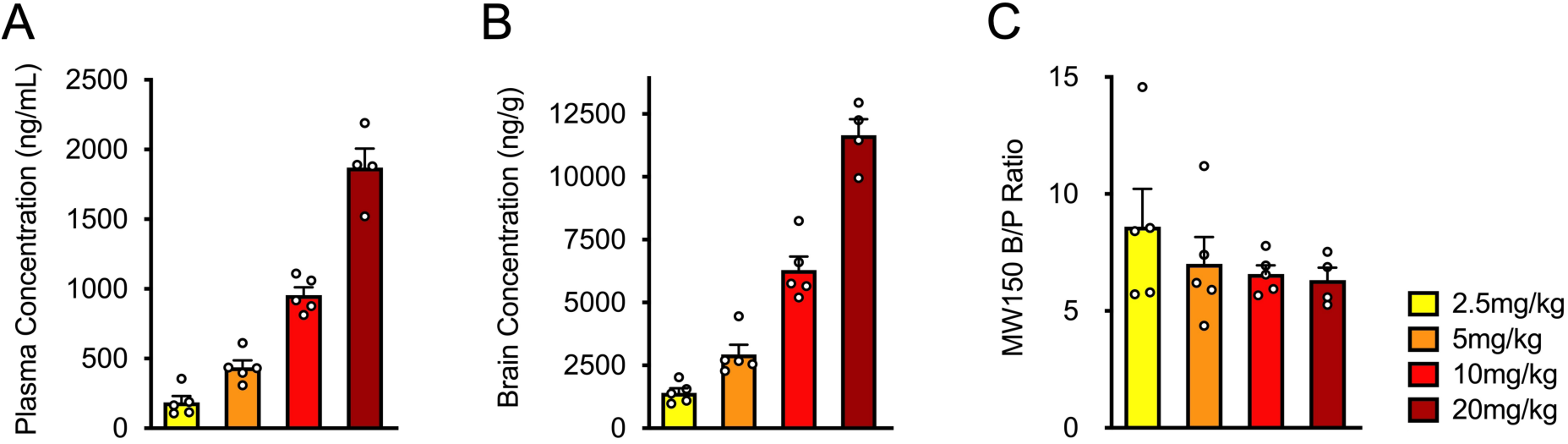
Analysis of MW150 biodistribution in plasma and brain of SMA mice. **(A-C)** MW150 concentration in plasma (A) and brain (B) and brain to plasma ratio (C) 3 hours after a single IP injection of the indicated doses MW150 in SMA mice at P10. Mean, SEM, and individual values from independent biological replicates are shown.

**Figure S2.**
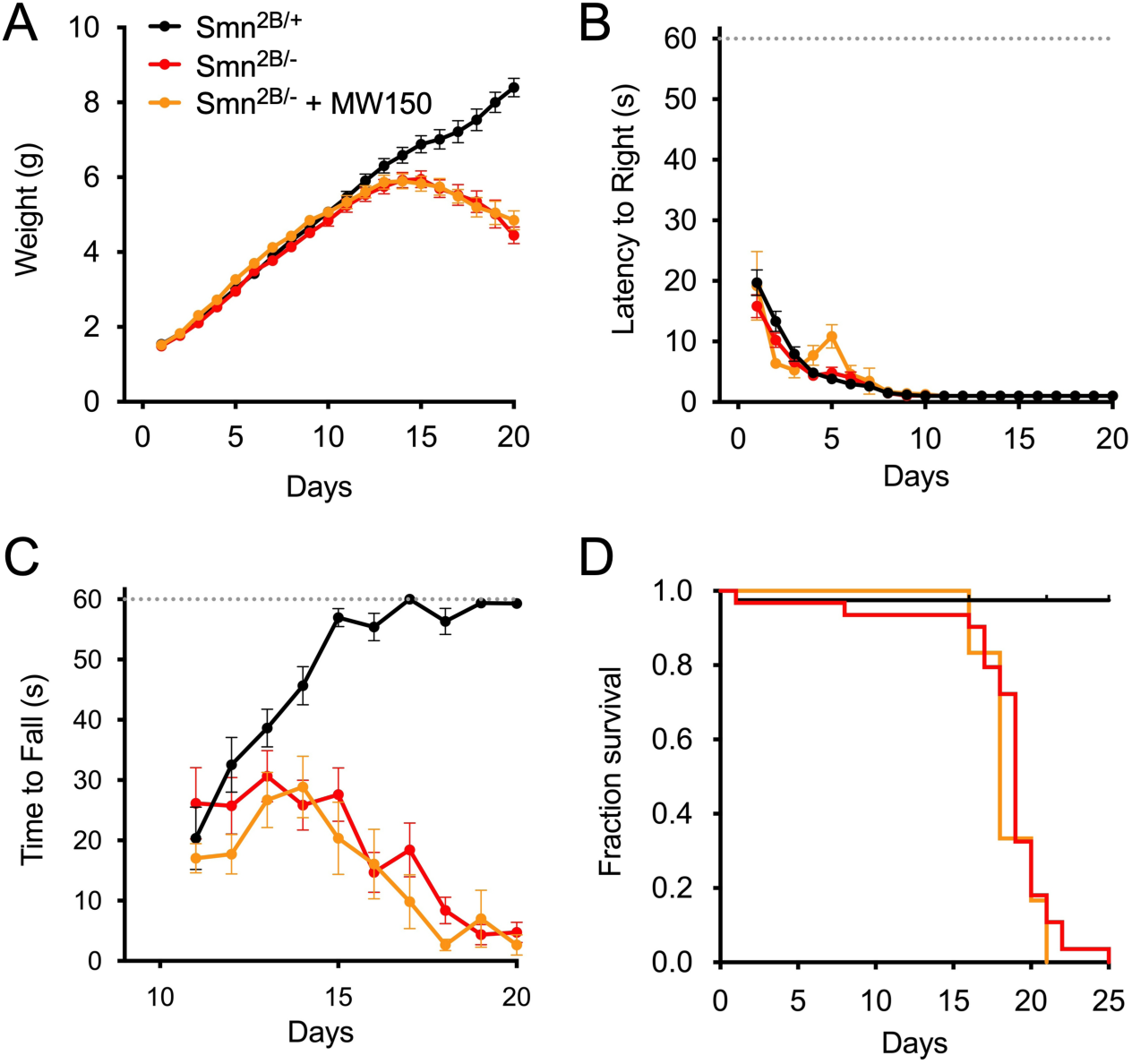
MW150 treatment does not improve the SMA phenotype in *Smn*^2B/-^ mice. (**A**) Body weight of control *Smn^2B/+^* (n=32) mice and *Smn^2B/-^* SMA mice either untreated (n=31) or treated daily with MW150 (5mg/kg) from P1 onward (n=12). Data represent mean and SEM. (**B**) Righting time from the same experimental groups shown in (A). Data represent mean and SEM. (**C**) Time to fall in the hindlimb suspension test from the same experimental groups shown in (A). Data represent mean and SEM. (**D**) Kaplan-Meier survival curves from the same experimental groups as in (A). The data for untreated *Smn^2B/+^* and *Smn^2B/-^* mice are from a previously published study (56).

**Supplementary Table 1.** List of antibodies used in this study.

| Name | Source | Cat # | Host | Application | Dilution |
| --- | --- | --- | --- | --- | --- |
| Phospho-p38 MAPK (Thr180/Tyr182) | Cell Signaling | 4511 | Rabbit | WB | 1:1,000 |
| p38 MAPK | Cell Signaling | 8690 | Rabbit | WB | 1:1,000 |
| SMN | BD | 610646 | Mouse | WB | 1:10,000 |
| α-Tubulin | Sigma-Aldrich | T6199 | Mouse | WB | 1:50,000 |
| β-Actin | Proteintech | 66009-1-Ig | Mouse | WB | 1:50,000 |
| GAPDH | Sigma-Aldrich | MAB374 | Mouse | WB | 1:50,000 |
| Peroxidase AffiniPure™ Goat Anti-Mouse | Jackson | 115-035-044 | Goat | WB | 1:10,000 |
| Peroxidase AffiniPure™ Goat Anti-Rabbit | Jackson | 111-035-003 | Goat | WB | 1:10,000 |
| ChAT | Sigma-Aldrich | AB144P | Goat | IHC | 1:100 |
| VGluT1 | Covance | Custom | Guinea Pig | IHC | 1:5,000 |
| Synaptophysin1 | Synaptic Systems | 101 308 | Guinea Pig | IHC | 1:500 |
| Neurofilament M | Sigma-Aldrich | AB1987 | Rabbit | IHC | 1:500 |
| Cy™3 AffiniPure™ Donkey Anti-Goat | Jackson | 705-165-147 | Donkey | IHC | 1:250 |
| Alexa Fluor® 488 AffiniPure™ Donkey Anti-Rabbit | Jackson | 711-545-152 | Donkey | IHC | 1:250 |
| Cy™5 AffiniPure™ Donkey Anti-Guinea Pig | Jackson | 706-175-148 | Donkey | IHC | 1:250 |

